# Antibacterial Effect of Oral Care Gel-Containing Hinokitiol and 4-isopropyl-3-methylphenol Against Intraoral Pathogenic Microorganisms

**DOI:** 10.1101/2023.03.07.531591

**Authors:** Hiroshi Ohara, Keita Odanaka, Miku Shiine, Masataka Hayasaka

## Abstract

**Objective:** Deterioration of oral hygiene is closely related to increase severity and mortality of corona virus disease-19 (COVID-19), and also contribute to the development of various diseases such as aspiration pneumonia or Alzheimer’s. Oral care is attracting high interest in Japan, which has entered a super-aging society. In this study, we aimed to investigate whether commercially available Hinora^®^ (HO) that oral care gels-containing hinokitiol and 4-isopropyl-3-methylphenol (IPMP) have biofilm formation inhibitory and antibacterial activities against various intraoral pathogen microorganisms.

**Method:** *Candida* spp., *Aggregatibacter actinomycetemcomitans, Staphylococcus aureus*, and *Pseudomonas aeruginosa* were selected during the study period, all which were analyzed using antimicrobial disc, microorganism turbidity, and crystal violet assays. In addition, the germ tube test using *C. albicans* was performed with a modification of Mackenzie’s method. Images for morphological observation of the germ tubes were acquired with an inverted microscope. For comparison between products, we used Refrecare^®^ (RC), which contains only hinokitiol (not containing IPMP).

**Results:** All the intraoral pathogenic microorganisms showed drug susceptibility against undiluted form HO and/or RC. In particular, HO was more effective at lower concentrations than RC. In the HO-added group, inhibition circles were observed in all bacteria except *P. aeruginosa* when added at a concentration of 0.5 g/mL or more. The optical density values at 590 nm (crystal violet) and/or 600 nm (microorganism turbidity) of all the fungi and bacteria were significantly lower when cultured in medium with HO. Inhibition of growth or biofilm formation was observed when HO was added at a concentration of 0.05 g/mL or higher. To investigate the action mechanism of HO, germ tube tests were performed in *C. albicans*. The results show that culturing *C. albicans* in soyabean-casein digest broth with HO (0.05 g/mL) significantly suppressed germ tube formation.

**Conclusions:** These data suggest that oral care gel-containing hinokitiol and IPMP has strong biofilm formation inhibitory, antifungal and antibacterial effects against *Candida* fungi and multiple intraoral pathogenic microorganisms. Therefore, it may be a promising treatment option for oral infections.

## Introduction

Oral infections such as oral candidiasis and periodontal disease is a series of polymicrobial infection that affect the oral mucosa and tooth root [1]. Deterioration of oral hygiene leads to intraorally colonization of fungi or bacteria, it has been reported to contribute to systemic diseases such as gingivitis, aspiration pneumonia, deep mycosis, or increased severity and mortality in the historical pandemic of corona virus disease-19 (COVID-19) [2–4]. It is suggested that aspiration pneumonia, which is often seen in the elderly, is caused by oral bacteria growing in the oral cavity. Oral care is attracting high interest in Japan, which has entered a super-aging society. Opportunistic infectious pathogenic microorganisms such as *Candida albicans* are known to easily form multi-species biofilms coaggregation on the surfaces of teeth and dentures [3, 5]. *Candida* spp. are facultative anaerobes and can grow in an environment where oxygen is sufficiently supplied, such as the oral surface and tooth surface, and in an environment where oxygen concentration is low, such as between teeth and periodontal pockets. *Candida albicans, C. glabrata, C. krusei, C. parapsilosis*, and *C. tropicalis* have been isolated from the oral cavity. *Candida albicans* is the most isolated species, but in the recent years, there have been reports of infection by other bacterial species. [6, 7] Proliferation of these fungi in the oral cavity can lead to superficial gingival inflammation, periodontitis, and deep-seated invasive gingival inflammation. [8]. Oral infection caused by *Candida* spp. is more likely to develop owing to immunodeficiency or decreased immunity owing to diabetes or human immunodeficiency virus infection. [9–11] In addition, unsanitary condition of oral has been reported to contribute to the development of Alzheimer’s disease and/or increased severity and mortality in the historical pandemic of severe acute respiratory syndrome coronavirus 2 (COVID-19). [12–17] Hence, periodic oral examinations and maintenance of oral hygiene are important.

To prevent exacerbation of oral hygiene, it is important to keep the oral cavity clean by brushing. In the treatment of oral infection, cleaning the oral cavity with oral care and antiseptic as well as treatment with antifungal drugs such as amphotericin B and miconazole is very important. However, *Candida* spp. are known to easily form biofilms on the surfaces of teeth and dentures, thereby reducing the effectiveness of brushing and antifungal agents. [18–20] The constituents of biofilms are polysaccharides and dead and live fungi. [21] Given that biofilm has high adhesion to the tooth surface, it is difficult to remove them all by the self-cleaning action of saliva or brushing. [22] In addition, given that the biofilm forms a membranous structure, the drugs found in mouthwashes do not penetrate the biofilm, reducing its effectiveness. [23, 24] Therefore, suppressing the formation of biofilms on the surfaces of teeth and dentures is very important for the prevention and treatment of oral infectious diseases.

Beta-thujaplicin (hinokitiol) is a tropolone-related compound extracted and purified from *Cupressaceae* plants such as *Chamaecyparis obtusa* and *Thujopsis dolabrata*. [25, 26] Hinokitiol has antibacterial effects and has been used in many daily necessities, foods, and cosmetics owing to its low toxicity to humans. [27–31] However, the biological activity of this compound is not yet fully understood and there are few reports on its antibacterial effects against orally pathogenic microorganisms. In this study, we aimed to investigate whether oral care gels-containing hinokitiol and 4-isopropyl-3-methylphenol (IPMP) (Hinora^®^: HO) or contains only hinokitiol (Refrecare^®^: RC) have biofilm formation inhibitory and antibacterial activities against fungi and bacteria that cause oral disease.

## Material and methods

### Strains and culture conditions

*Candida albicans (C. albicans*) (ATCC10231), *A. actinomycetemcomitans* (ATCC29522), *P. aeruginosa* (ATCC9027), *S. aureus* (ATCC6538), *C. glabrata* (NBRC0622), *C. krusei* (NBRC1395), *C. parapsilosis* (NBRC1396), and *C. tropicalis* (NBRC1400) were purchased from the American Type Culture Collection (ATCC, Virginia, USA) or Biological Resource Center, National Institute of Technology and Evaluation (NBRC, Chiba, Japan). All bacteria and fungi, except *A. actinomycetemcomitans*, were cultured under aerobic conditions, and *A. actinomycetemcomitans* was cultured under anaerobic conditions (5 % CO_2_, 0 % O_2_), using Anaero Pouch Kenki (Mitsubishi Gas Chemicals Co., Ltd., Tokyo, Japan).

In the culture using soyabean-casein digest (SCD) agar medium (Nissui pharmaceutical Co., Ltd., Tokyo, Japan), each fungus and bacterium were cultured at 37 °C for 24–48 h. For the culture using SCD broth (Merck KGaA, Darmstadt, Germany) with and without HO (EN Otsuka Pharmaceutical Co., Ltd., Iwate, Japan) and/or RC (Bean Stalk Snow Co., Ltd., Tokyo, Japan) (HO concentration: 0.05 g/mL, 0.125 g/mL; RC concentration: 0.05 g/mL, 0.125 g/mL), the obtained fungal and bacterial suspensions were dispensed onto 96-well polystyrene plates at 200 μL/well and cultured at 37 °C for 24–48 h. For all experiments, each fungus was adjusted to a 0.1–0.4 optical density at 600 nm (equivalent to 1 × 10^6^ colony-forming units (CFU) /mL) and inoculated at 1 × 10^6^ CFU/mL, and each bacterium was adjusted to a 0.1–0.3 optical density at 600 nm (equivalent to 1 × 10^8^ CFU/mL) and inoculated at 1 × 10^8^ CFU/mL.

### Antimicrobial disc diffusion assay

Each fungus and bacterium were inoculated into saline solutions to a concentration of 1 × 10^6^ or 1 × 10^8^ CFU/mL and spread evenly on SCD agar medium. A sterilized paper disc with a diameter of 8 mm (ADVANTEC Toyo, Co., Ltd., Tokyo, Japan) coated with oral care gel-containing medium was placed on the SCD agar medium, and the fungus or bacterium was cultured at 37°C for 24 h.

Paper discs were coated with 80 μL of SCD broth and SCD broth containing HO (0.1, 0.5, and 5 g/mL and undiluted form) or RC (0.1, 0.5, and 5 g/mL and undiluted form). A paper disc coated with oral care gel-containing medium was used after 2–3 h of air drying in a clean work station.

### Microorganism turbidity and crystal violet assays

As described above, each fungus and bacterium were cultured at 37 °C for 24–48 h in SCD broth and SCD broth containing HO (0.05 g/mL, 0.125 g/mL) or RC (0.05 g/mL, 0.125 g/mL). First, the absorbance of cultures was measured at optical density (OD) 600 nm on a SpectraMax190 microplate reader (Molecular Devices Co., Ltd., Tokyo, Japan) to determine the turbidity (i.e. level of proliferation of each fungus and bacterium). Next, the culture was removed, and each well was washed twice with saline. A crystal violet aqueous solution (0.1 % w/v) was added to each well (200 μL/well), and the mixture was allowed to stand at room temperature (approximately 25 °C) for 30 min for staining. After staining, the crystal violet aqueous solution was removed, and each well was washed twice with saline. Ethanol was added to the well (200 μL/well), and the mixture was allowed to stand at room temperature (approximately 25 °C) for 15 min. Then, the crystal violet stain solution was extracted from each fungus or bacterium with ethanol, and the optical density was measured at OD 590 nm on a microplate reader to detect the biofilm in the 96-well plate.

### Germ tube test

The germ tube test was a modification of Mackenzie’s method. [32] *Candida albicans* was inoculated into SCD broth or SCD broth with HO (0.05 g/mL) to a concentration of 1 × 10^6^ CFU/mL. The fungal suspension was applied in a 6-cm dish (3 mL/dish) and cultured at 37 °C for 3 h. Images for morphological observation of the germ tubes were acquired with an Olympus CKX41 camera (Olympus Co., Ltd., Tokyo, Japan).

### Statistical analysis

The statistical significance of the differences in the ratio of biofilm formation between non-treated bacterial pathogens and HO- and RC-treated bacterial pathogens was examined using Dunnett’s test. The statistical significance level was considered to be statistically significant when *P* < 0.05.

## Results

The inhibitory zone formed by the undiluted solution of HO and RC for each *Candida* fungus was larger than that of the control group (Fig. 1). The inhibition zone was larger for HO than for RC, with diameters of 24 mm for *C. albicans* (Fig. 1A), 25 mm for *C. glabrata* (Fig. 1B), 25 mm for *C. krusei* (Fig. 1C), 24 mm for *C. parapsilosis* (Fig. 1D), and 23 mm for *C. tropicalis* (Fig. 1E). In addition, all oral infection-causing bacteria used in the experiments formed inhibitory zones, and the maximum zone of inhibition was 31 mm of *S. aureus* in the HO-added group. Table 1 shows the details of the inhibition zone diameter. In the HO-added group, inhibition circles were observed in all bacteria except *P. aeruginosa* at a concentration of 0.5 g/mL or higher. In the RC-added group, inhibition circles were observed in all bacteria except *C. krusei* at a concentration of 5 g/mL or higher (Table 1).

**Fig 1.**
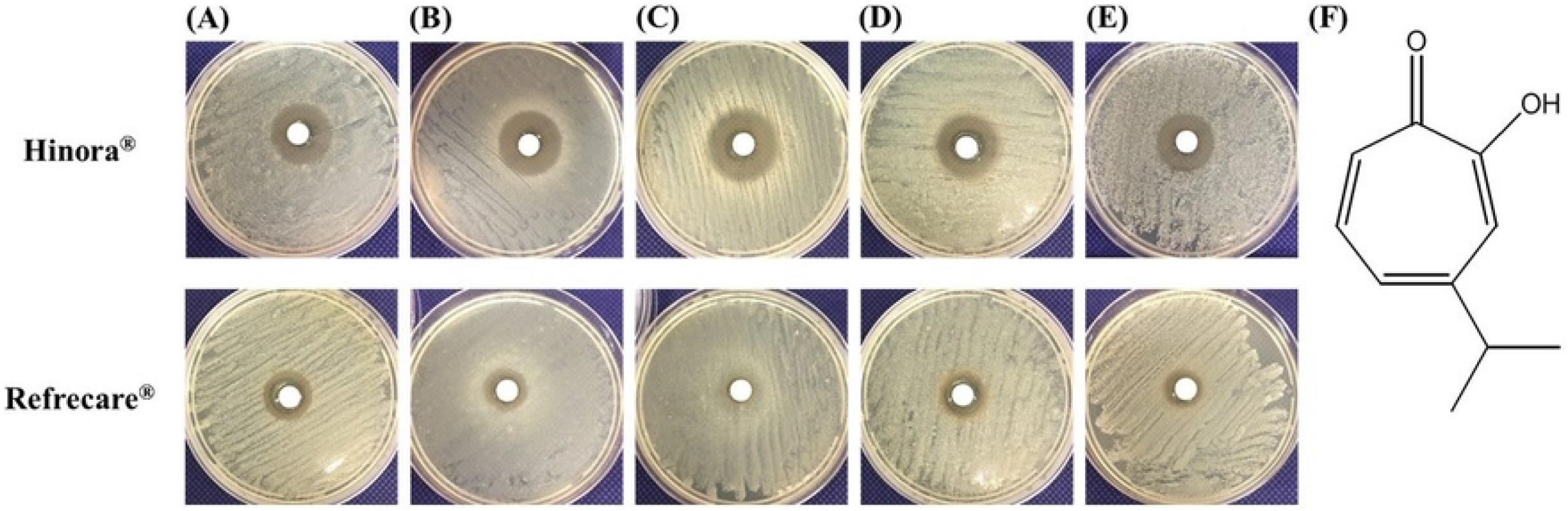

We next cultured various bacteria in SCD broth containing HO or RC (final 0.05, 0.25 g/mL) for 24 h, and bacterial growth was measured using spectrophotometry by absorbance at OD 600 nm. The value of OD 600 nm was significantly lower when cultured in SCD broth containing HO or RC compared to the control (Fig. 2). These growth suppression effects were observed when the concentration of HO or RC was 0.05 g/mL or higher.

**Fig 2.**
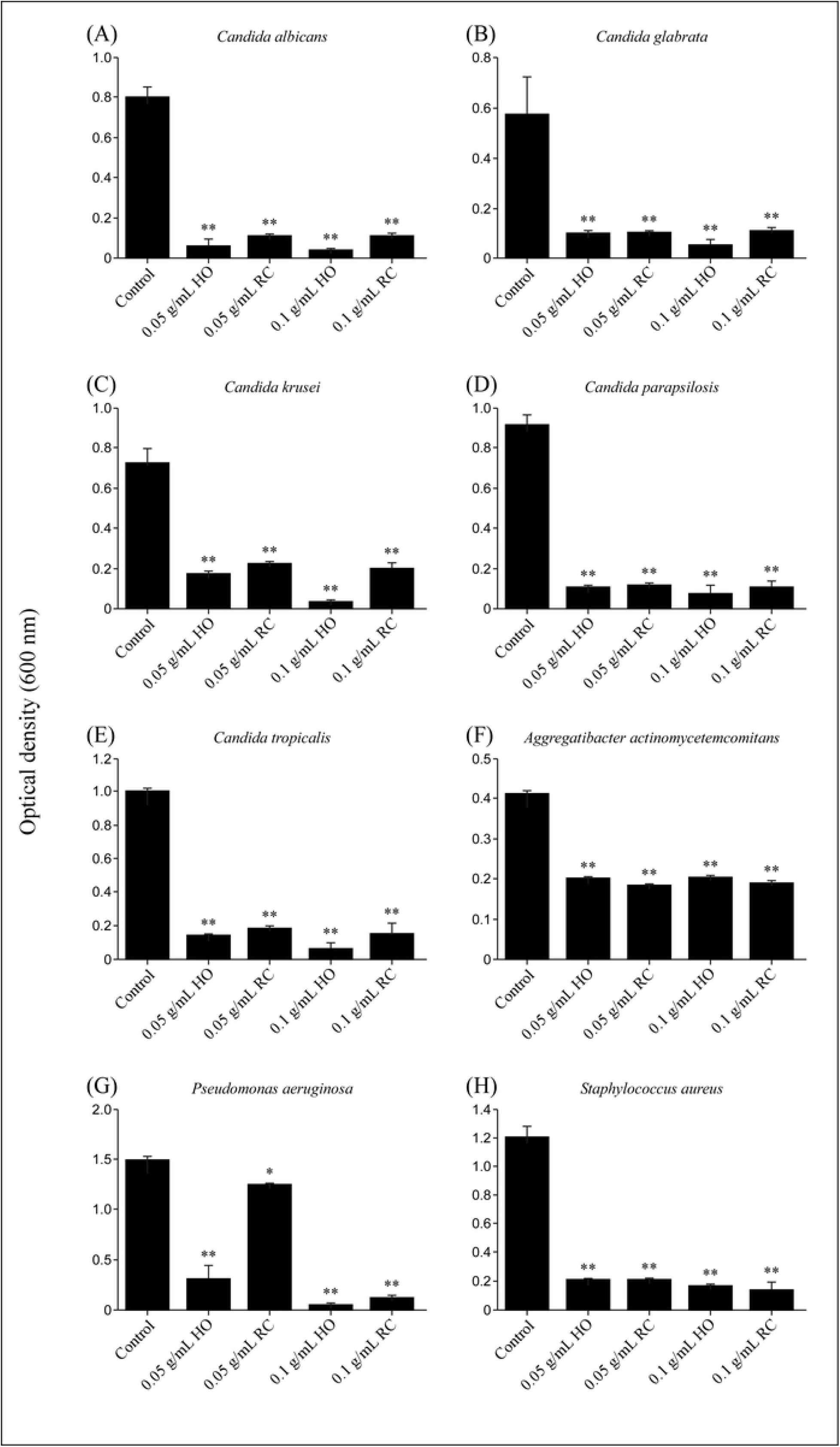

Subsequently, we investigated whether these oral care gels had biofilm formation inhibitory activity against various bacteria using crystal violet assays. The addition of HO inhibited biofilm formation of all the oral infection-causing bacteria used (Fig. 3). The inhibitory effect of HO was observed at 0.05 g/mL or higher. On the other hand, RC showed a biofilm formation inhibitory effect against bacteria other than *A. actinomycetemcomitans*. The inhibitory effect of RC was observed at 0.05 g/mL or higher.

**Fig 3.**
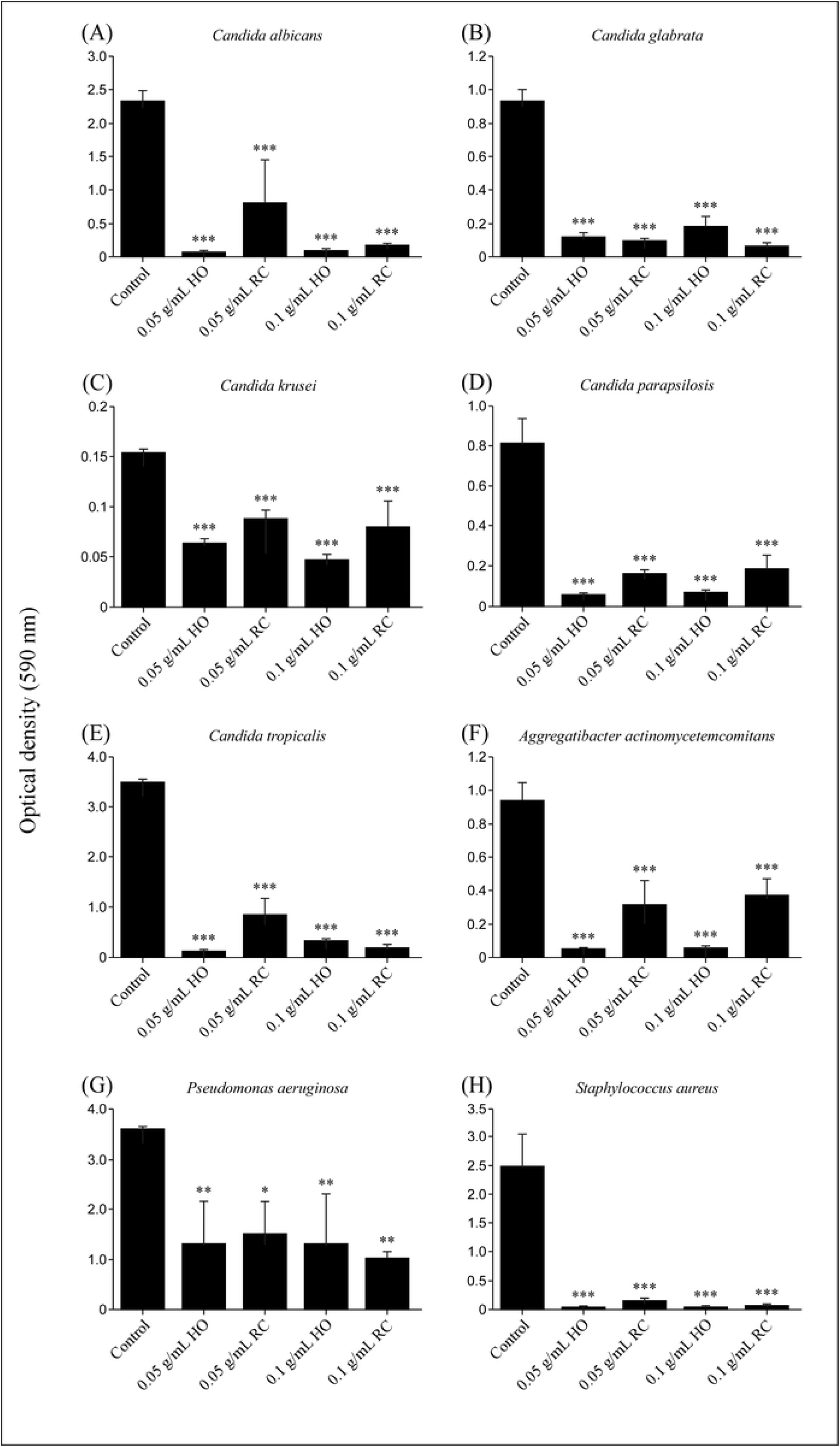

Finally, we conducted a germ tube test of *C. albicans* on a HO-containing SCD broth to investigate the mechanism of action of HO, which had an excellent ability to inhibit biofilm formation. The results show that the addition of HO significantly suppressed the formation of germ tubes in *C. albicans* (Fig. 4). Germ tube formation was inhibited even at a concentration as low as 0.05 g/mL of HO.

**Fig 4.**
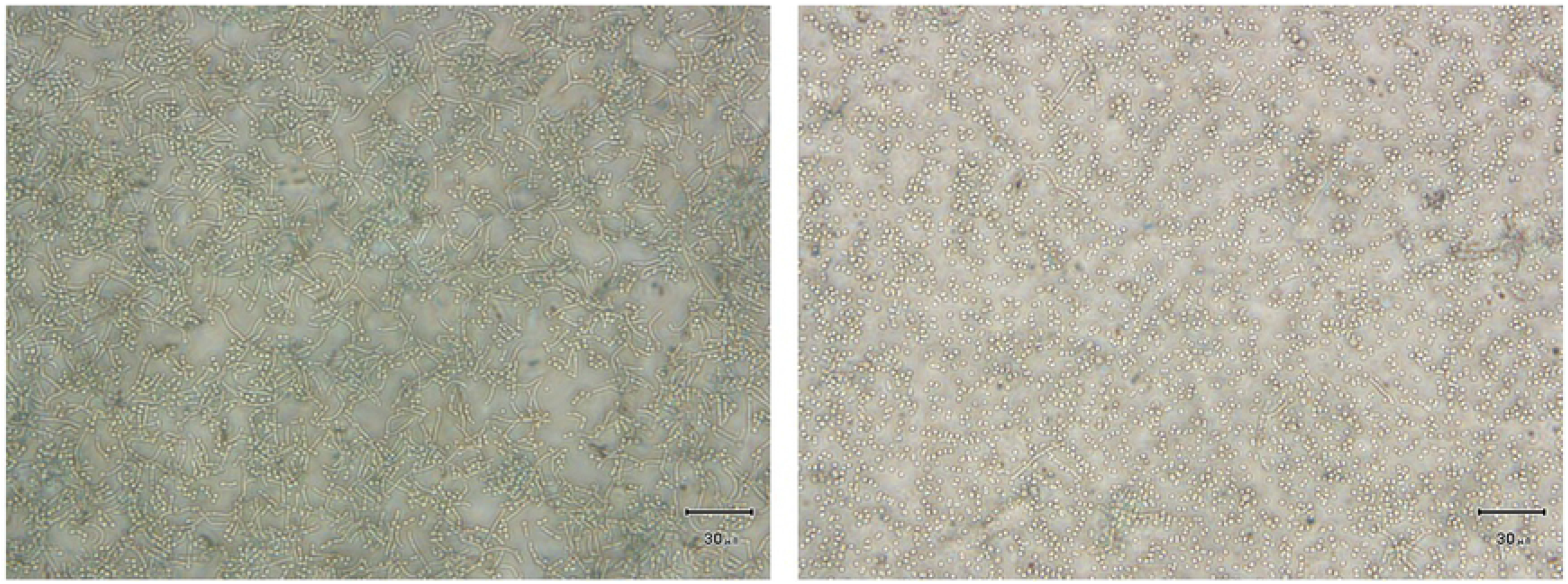

## Discussion

Whether the oral care gel Hinora^®^ (HO) and Refrecare^®^ (RC) showed antifungal and antibacterial activity was examined using the disc method. In addition to *Candida* spp. (*C. albicans* (ATCC10231), *C. glabrata* (NBRC0622), *C. krusei* (NBRC1395), *C. parapsilosis* (NBRC1396), and *C. tropicalis* (NBRC1400)), we also investigated *A. actinomycetemcomitans* (ATCC 29522), *P. aeruginosa* (ATCC9027), and *S. aureus* (ATCC6538), which are often isolated from periodontal disease patients. The onset of oral infection caused by these fungi and bacteria is common in people with reduced immunity, such as patients with acquired immunodeficiency syndrome. [1, 19, 33]

Figure 1 shows the growth inhibition zones of *Candida* spp. For all fungi and bacteria that cause oral infectious diseases, growth-inhibiting circles were formed around the disc containing HO or RC (Table 1). In the HO-added group, inhibition circles were observed in all bacteria except *P. aeruginosa* when added at a concentration of 0.5 g/mL or more. In the RC-added group, inhibition circles were observed in all bacteria except *C. krusei* when added at a concentration of 5 g/mL or more (Table 1).

In addition, various fungi and bacteria described above were cultured in SCD broth with and without HO or RC for 24 h, and the optical density of the culture was measured at 600 nm on a microplate reader to detect the turbidity (indicative of the level of proliferation of each fungus and bacterium). As a result, the optical density values at 600 nm of all the cultures were significantly lower when cultured in medium with HO or RC as compared to those of cultures without HO or RC (Fig. 2). Growth inhibition was observed when HO or RC was added at a concentration of 0.05 g/mL or higher (Fig. 2). These results indicate that HO and RC showed growth inhibitory effects on the microorganisms used in this study.

Furthermore, the results of the disc diffusion assay and culture in SCD liquid medium showed that HO tended to have higher antibacterial activity than that of RC. Hinokitiol content was twice as high in HO as compared to that in RC (HO: 0.1 %; RC: 0.05 %). Moreover, 0.02 % 4-isopropyl-3-methylphenol (IPMP) was present in HO but not in RC. Komaki N et al reported that the growth inhibition of *C*. *albicans* by hinokitiol occurs via inhibition of the *RAS1* signaling pathway, one of the cAMP-dependent protein kinases. [34] IPMP is typical of preservatives and bactericidal ingredients in cosmetics and is used in many oral care products. [35, 36] These facts are consistent with the observation that HO tended to have higher antibacterial activity than RC.

A crystal violet assay was performed to determine whether these oral care gels had biofilm formation inhibitory activity. Oral infection-causing microorganisms such as *Candida* spp. form biofilm on the tooth surface, resisting chemical and physical treatments such as antibacterial mouthwash and brushing. [22–24] In the oral cavity, the explosive growth of pathogens causes oral infection to worsen and become intractable. Therefore, it is very important to suppress biofilm formation on the tooth surface for the prevention and treatment of oral infectious diseases. We investigated whether oral care gels HO and RC could inhibit biofilm formation caused by oral infection-causing microorganisms. As a result, the addition of HO inhibited the formation of biofilms of all the oral infection-causing bacteria studied (Fig. 3). Addition of HO at concentrations above 0.05 g/mL strongly inhibited biofilm formation in all the fungi. RC showed biofilm formation inhibitory effects on all the microorganisms examined, except *A. actinomycetemcomitans*. When the RC concentration was 0.05 g/mL or higher, the biofilm formation inhibitory effect of microorganisms was observed. The biofilm formation inhibitory effects of HO and RC in *Candida* spp. were estimated to be comparable. However, given that HO showed a biofilm formation inhibitory effect on more bacteria than RC, it is inferred that the use of HO could be more practical than RC.

Biofilm formation occurs during the initial stage of adhesion to the tooth surface. Microorganisms grow in the oral cavity, attach to the tooth surface, and accumulate, forming a membrane-like structure. Biofilms are composed of polysaccharides, multiple ions, and microorganisms. The growth rate of microorganisms in biofilms is slow, but not all bacteria inside the biofilm die. In contrast, microorganisms on the surface of biofilms can spread the infectious region as they move and multiply in search of other habitats as needed. [21, 37, 38] Germ tube formation is one of the processes involved in adhesion, which is the first step in the biofilm formation of *Candida* spp.. [22, 39] The formation of germ tubes is known to be promoted by stimulation with proteins in serum at 37 °C and is easily formed even in SCD broth. [32, 40] In this study, to investigate the mechanism of action of HO, *C. albicans* was cultured in HO-containing SCD broth, and its effect on germ tube formation was investigated. The results showed that culturing *C. albicans* in SCD broth with HO (0.05 g/mL) suppressed germ tube formation (Fig. 4). Kim DJ et al has demonstrated that hinokitiol suppresses the expression of genes involved in *C. albicans* adhesion process (*HWP1* and *ALS3*) and hyphal formation (*UME6*) and/or maintenance (*HGC1*), and claims that these results are mechanisms that inhibit biofilm formation. [41]

HO is presumed to suppress biofilm formation by inhibiting the adhesion of *C. albicans* to the tooth surface. In the future, it will be necessary to investigate in detail the mechanism of HO and its mechanism of action in other microorganisms as well. In addition, given that these oral care gels exerted a biofilm formation inhibitory effect and an antibacterial effect against *Candida* spp. at a low concentration (0.05 g/mL), the effects of the levels used for oral care should be investigated. Moreover, the optimal use of these gels, that is, whether to use these when brushing teeth, to gargle these with water after brushing teeth, and so on, should be considered in future studies.

## Conclusion

In conclusion, oral care gel-containing hinokitiol were found to have biofilm formation inhibitory and antibacterial activities against the various intraoral pathogenic microorganisms used in this study. In particular, oral care gel-containing hinokitiol and 4-isopropyl-3-methylphenol (IPMP) strongly inhibited biofilm formation and had antibacterial effects against multiple oral infection-causing bacteria (*A. actinomycetemcomitans* [ATCC29522], *P. aeruginosa* [ATCC9027], and *S. aureus* [ATCC6538]) other than *Candida* spp. These results suggest that oral care gel-containing hinokitiol are considered very effective in the prevention and treatment of oral infections such as oral candidiasis or periodontal disease. In addition, oral care gel-containing hinokitiol and IPMP may be particularly useful for assisting the prevention and treatment of *Candida* fungi and multiple oral infectious microorganisms.

## Acknowledgments

We are also grateful to Kenichi Ogasawara and Ryosuke Kawawaki of EN Otsuka Pharmaceutical Co., Ltd. for their advice in promoting this research.

